# Tmem178 negatively regulates IL-1β production through inhibition of the NLRP3 inflammasome

**DOI:** 10.1101/2023.03.07.531385

**Authors:** Kunjan Khanna, Hui Yan, Muneshwar Mehra, Nidhi Rohatgi, Gabriel Mbalaviele, Roberta Faccio

**Affiliations:** Department of Orthopedics, Washington University in St Louis, MO, USA; Key Laboratory for Animal Disease-Resistance Nutrition of China Ministry of Education, Ministry of Agriculture and Rural Affairs and Sichuan Province, Animal Nutrition Institute, Sichuan Agricultural University, Chengdu 611130, China; Department of Neuroscience, Washington University in St Louis, MO, USA; Department of Pathology and Immunology, Washington University in St Louis, MO, USA; Department of Medicine, Washington University in St Louis, MO, USA; Shriners Hospital for Children, St Louis, MO, USA

**Keywords:** sJIA, NLRP3 inflammasome, IL-1β, Ca2+, Tmem178, Stim1

## Abstract

**Objective:** Inflammasomes modulate the release of bioactive IL-1β. Excessive IL-1β levels are detected in patients with systemic juvenile idiopathic arthritis (sJIA) and cytokine storm syndrome (CSS) with mutated and unmutated inflammasome components, raising questions on the mechanisms of IL-1β regulation in these disorders.

**Methods:** To investigate how the NLRP3 inflammasome is modulated in sJIA, we focused on Tmem178, a negative regulator of calcium levels in macrophages, and measured IL-1β and caspase-1 activation in wild-type (WT) and *Tmem178*^*-/-*^ macrophages following calcium chelators, silencing of Stim1, a component of store-operated calcium entry (SOCE), or by expressing a Tmem178 mutant lacking Stim1 binding site. Mitochondrial function in both genotypes was assessed by measuring oxidative respiration, mitochondrial reactive oxygen species (mtROS), and mitochondrial damage. CSS development was analyzed in *Perforin*^*-/-*^*/Tmem178*^*-/-*^ mice infected with LCMV in which inflammasome or IL-1 signaling was pharmacologically inhibited. Human *TMEM178* and *IL-1B* transcripts were analyzed in a dataset of peripheral blood monocytes from healthy controls and active sJIA patients.

**Results:** *TMEM178* levels are reduced in monocytes from sJIA patients while IL-1B show increased levels. Accordingly, *Tmem178*^*-/-*^ macrophages produce elevated IL-1β compared to WT cells. The elevated intracellular calcium levels following SOCE activation in *Tmem178*^*-/-*^ macrophages induce mitochondrial damage, release mtROS, and ultimately, promote NLRP3 inflammasome activation. *In vivo*, inhibition of inflammasome or IL-1 neutralization prolongs *Tmem178*^*-/-*^ mouse survival to LCMV-induced CSS.

**Conclusion:** Downregulation of Tmem178 levels may represent a new biomarker to identify sJIA/CSS patients that could benefit from receiving drugs targeting inflammasome signaling.

## Introduction

Systemic juvenile idiopathic arthritis (sJIA) is a serious clinical complication of systemic inflammatory disorders characterized by arthritis and fever (1) (2). Almost 300,000 children in U.S.A. are currently fighting some form of arthritis, which is more than the number of patients for cystic fibrosis, leukemia and juvenile diabetes combined together. About 3-10% of sJIA patients develop a life-threatening complication named macrophage activation syndrome (MAS) or cytokine storm syndrome (CSS), where uncontrolled T cell and macrophage activation result in an exaggerated systemic inflammatory response (3). Although the causes of MAS/CSS are unknown, this complication is also observed in critically ill, COVID19 patients (4), cancer patients receiving CAR T-cell immunotherapies (5), and patients with Kawasaki disease (6). Thus, understanding the driver/s of this deadly inflammatory storm can have important therapeutic implications for a variety of conditions.

Studies using animal models as well as samples from sJIA patients developing CSS have shown elevated levels of pro-inflammatory cytokines in particular IL-1β and IL-18 (7) (8) (9). IL-1β and IL-18 are modulated by multimolecular complexes, named inflammasomes, sensing microbe-derived pathogen-associated molecular patterns (PAMPs) and danger-associated molecular patterns (DAMPs) (10). Activation of caspases, caspase-1 in particular, by the inflammasomes allows the cleavage of the immature pro-inflammatory cytokines pro-IL-1β and pro-IL-18 into their active forms and their subsequent release (11).

Whole exome sequencing in sJIA patients developing recurrent CSS identified a variant in the gene *CASP1*, which encodes for caspase-1, leading to gain-of-function of inflammasomes (12). An activating-NLRC4 mutation was further shown to cause autoinflammation with recurrent CSS (13). Similarly, NLRP3 inflammasome overactivation has been observed in CSS patients infected by SARS-CoV-2 (14). However, inflammasome mutations or variants in *CASP1* are extremely rare, underscoring the need to understand how inflammasomes are activated during CSS and whether inflammasome-mediated inflammatory responses are drivers of disease progression.

We recently identified Tmem178, a transmembrane protein in the endoplasmic reticulum, as an important negative modulator of inflammatory cytokine production in macrophages through regulation of calcium fluxes (15). Macrophages lacking Tmem178 develop severe inflammatory responses in two mouse models of MAS/CSS, one induced by repeated TLR9 stimulation in response to CpG administration and a second driven by lymphocytic choriomeningitis virus (LCMV) infection in the perforin null background (16). The drivers of disease severity in *Tmem178*^*-/-*^ mice remain to be defined.

In this study, we demonstrate that loss of *Tmem178* induces NLRP3 inflammasome activation and IL-1β release via increased Ca^2+^ levels through Stim1, a component of the store operated calcium entry (SOCE), and mitochondrial damage. *Tmem178*^*-/-*^ mice show prolonged survival to LCMV-induced CSS following administration of an inflammasome signaling inhibitor or IL-1β targeting. Importantly, *Tmem178* levels are reduced in a sJIA patient dataset of human monocytes and negatively correlate with IL-1β expression. Our data identifies Tmem178 as a novel negative modulator of IL-1β production.

## Materials and Methods

Details on materials and procedures for *in vitro* and *in vivo* experiments used in the present study are described in Supplementary Materials.

### Mice

WT, *Tmem178*^*-/-*^, *Nlrp3*^*-/-*^, *Prf*^*-/-*^, and correspondent double knockout mice were used for the experiments. *Tmem178*^*-/-*^ mice (Strain B6;129S5-Tmem178tm1Lex/Mmucd) were generated by the trans-NIH Knock-Out Mouse Project (KOMP) and purchased from the KOMP Repository at University of California Davis (https://www.komp.org) (stock no. 032664-UCD) and crossed by speed congenics to obtain pure C57/B6 background. C57/B6 *WT, Nlrp3*^*-/-*^and *Prf*^*-/-*^ *m*ice were purchased from Jackson Laboratory (Bar Harbor, ME). *Tmem178*^*-/-*^ mice were crossed with *Nlrp3*^*-/-*^ or *Prf*^*-/-*^ mice to generate *Tmem178*^*+/-*^*;Nlrp3*^*+/-*^ or *Tmem178*^*+/-*^*;Prf*^*+/−*^ (double heterozygous, F1) mice, which were, in turn, inter-crossed to generate F2 homozygous breeders (*Tmem178*^*-/-*^*;Nlrp3*^*-/-*^ or Tmem178^*-/-*^*;Prf*^*-/-*^). F3 mice were used for all experiments. All mice were on C57/BL6 background and housed *ad libitum* with free access to feed and water. Female and male animals were used for all experiments since no gender differences were noted. When different groups were compared, the same numbers of males and/or females used in each group. All experiments were approved by Washington University School of Medicine animal care and use committee.

### *In vivo* animal models

For LPS-induced inflammation, LPS (5 or 10mg/kg) from Escherichia coli 0111:B4 (Sigma) was injected intraperitoneally into 6-8 week-old male mice. Animal were euthanized after 6 or 16 hours following LPS injection to collect blood, peritoneal fluid and liver samples for analysis of IL-1β levels. For the CpG-induced CSS model, 50μg CpG 1826 (IDT) was injected intraperitoneally into 6-8 week-old male mice every 2 days. On day 9, the mice were euthanized to collect liver samples to analyze IL-1β levels. For the LCMV-induced CSS model, 2×10^5^ plaque-forming units (PFUs) of LCMV-Armstrong were administered intraperitoneally in 8-12 week-old female *Prf*^*-/-*^ or *Prf*^*-/-/*^:*Tmem178*^*-/-*^ double knockout mice and animals were monitored daily to record survival.

### Statistical analysis

Data is represented as mean +/- SEM using GraphPad Prism 9. Statistical significance was determined by unpaired Student’s *t*-test or in experiment with multiple comparisons by the two-way ANOVA in conjunction with Sidak’s multiple comparison test, as indicated. For human dataset analysis, significance was determined by Wald test using DESeq2. For survival studies, the significance was determined by log-rank (Mantel-Cox) test. *P* value <0.05 is set as statistically significant.

## Results

### Tmem178 deficiency increases IL-1β production and NLRP3 inflammasome activation

To evaluate IL-1β levels in *Tmem178*^*-/-*^ mice under inflammatory conditions, we injected a sublethal dose of the TLR4 ligand LPS (5mg/kg) or delivered the TLR9 ligand CpG every other day for 9 days into 6-8 weeks old, sex matched WT and *Tmem178*^*-/-*^ animals. A significant increase in IL-1β was detected 16 hours after LPS stimulation in the serum, peritoneal fluid, and liver lysate of *Tmem178*^*-/-*^ mice compared to WT (Fig.1A). IL-1β levels were also increased in liver lysates of *Tmem178*^*-/-*^ mice following CpG stimulation, although remained undetectable in the serum and peritoneal fluid (Fig. 1B and data not shown).

**Figure 1.**
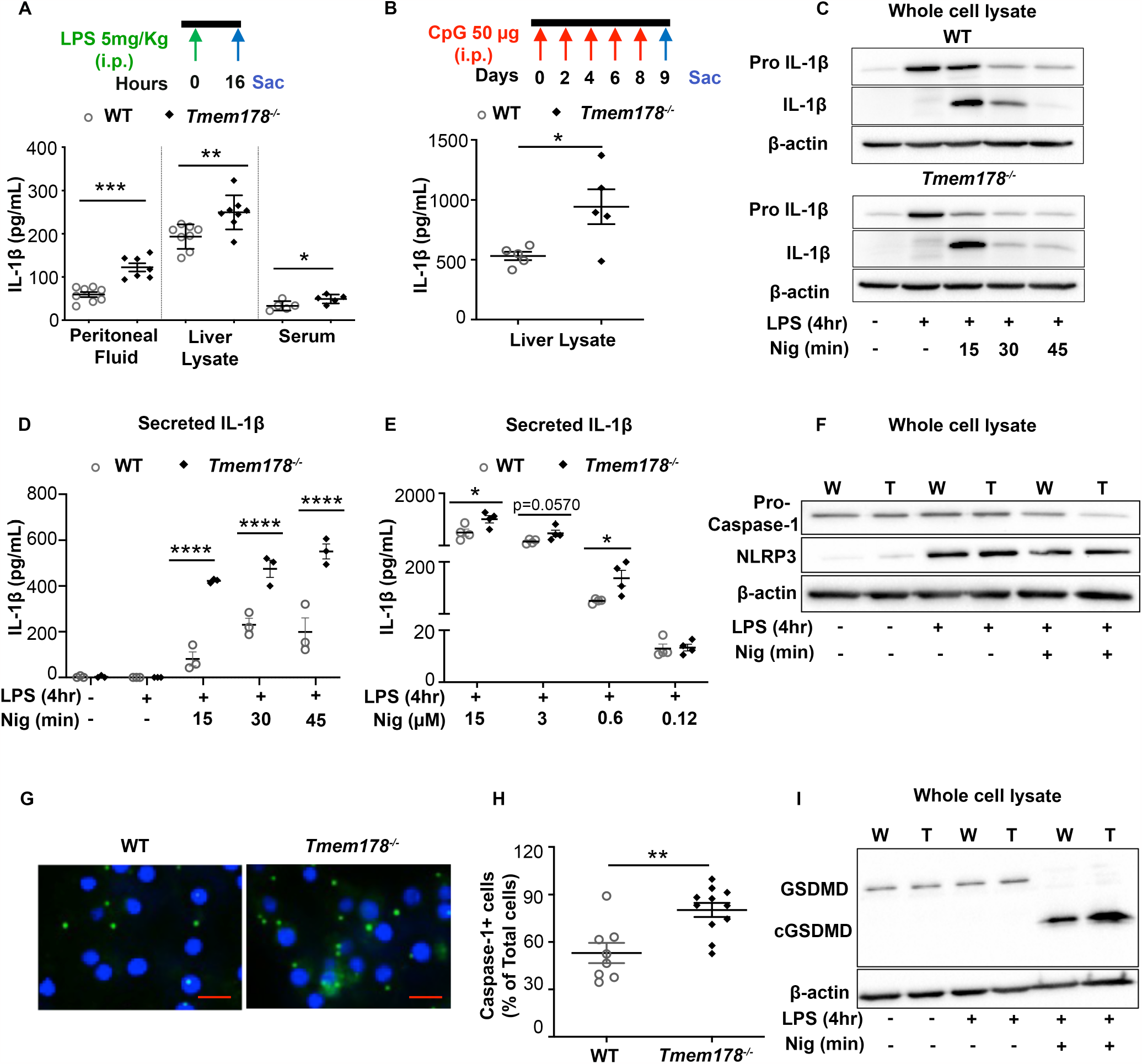
*Tmem178* deficiency increases IL-1β production. IL-1β levels in WT or *Tmem178*^*-/-*^ mice challenged with (A) 5mg/kg LPS i.p. (16 hours) in peritoneal fluid (n = 7-8/group), liver lysate (n = 8/group) and serum (n = 5/group) or (B) 50μg CpG i.p. (8 days) in liver lysates (n = 5/group). WT and *Tmem178*^*-/-*^ BMDMS were primed with 100ng/ml LPS for 4hr followed by nigericin for indicated time/dose and subjected to (C) western blotting to detect pro-IL-1β and mature IL-1β protein levels, (D-E) ELISA to measure IL-1β release in the culture supernatants (n = 3-4/group), (F) pro-caspase-1 and NLRP3 protein analysis by immunoblotting, (G) fluorescence imaging representing active caspase-1 (green) and DAPI staining (blue) (Scale=10μm), (H) quantification of percentage of cells with active caspase-1 in WT (n = 8) and *Tmem178*^*-/-*^ (n = 11) cultures, (I) GSDMD and cleaved GSDMD protein analysis in WT (W) and *Tmem178*^*-/-*^ (T) lysates. Data are expressed as mean ± SEM. Immunoblots are representative of three independent experiments and β-actin was used as loading control. Statistical significance is determined by two-tailed t-test (A, B, E, H) or two-way ANOVA for multiple comparisons (D). \**P* < 0.05, \*\**P* < 0.01, \*\*\**P* < 0.001 and \*\*\*\**P* < 0.0001.

Synthesis and release of IL-1β is a two-step process (10). In the first or priming step, activation of NF-κB induces expression of IL-1β transcripts and pro-IL-1β protein. In the second step, activation of the inflammasome induces the cleavage of pro-IL-1β into the mature active form and its release outside of the cell. Similar to osteoclasts (15), we did not detect differences in the phosphorylation and degradation of IκB-α, a readout for NF-κB activation (20), between WT and *Tmem178*^*-/-*^ bone marrow derived macrophages (BMDMs) stimulated with LPS (Supplemental Fig. 1A-C). However, *pro-IL-1β* transcripts were significantly higher in the *Tmem178*^*-/-*^ cell cultures compared with WT, while cytoplasmic pro-IL-1β protein levels were similarly increased by LPS in both genotypes (Supplemental Fig. 1D-E).

NLRP3 polymorphisms are associated with the pathophysiology of sJIA (21), and NLRP3 inflammasome overactivation is considered a trigger for development of CSS after SARS-CoV-2 infection (22). To determine whether Tmem178 deficiency increases IL-1β release by controlling the NLRP3 inflammasome, WT and *Tmem178*^*-/-*^ BMDMs were primed with LPS for 4 hours followed by stimulation with the NLRP3 inflammasome activator, nigericin. Intracellular pro-IL-1β levels were reduced at a faster rate in *Tmem178*^*-/-*^ cells, with a concomitant increase in IL-1β release compared with WT (Fig. 1C-D, Supplemental Fig. 1F-G). Furthermore, a dose-response to different concentrations of nigericin showed higher IL-1β levels in *Tmem178*^*-/-*^ compared to WT cultures (Fig. 1E).

Active caspase-1 cleaves pro-IL-1β into its mature form (23). While both genotypes expressed similar levels of pro-caspase-1 protein at baseline (Fig. 1F, Supplemental Fig. 1I), treatment with LPS and nigericin led to a slightly faster reduction of pro-caspase-1 in *Tmem178*^*-/-*^ BMDM (Fig. 1F). LPS induced NLRP3 expression as previously (24). To test the hypothesis that the activated state of the NLRP3 inflammasome is higher in knock-out compared to control cells, we quantified the percentage of cells with activated caspase-1, which binds the FLICA™ FAM-YVAD-FMK probe (25). We observed a higher number of cells with active caspase-1 in the *Tmem178*^*-/-*^ cultures compared with WT (Fig. 1G, H). By contrast, the levels of caspase-8, involved in inflammasome independent IL-1β release, were similar in WT and knock-out cells (not shown). Cleavage of gasdermin-D (GSDMD) is dependent on caspase-1 activation in response to NLRP3 inflammasome activators. We observed higher levels of cleaved GSDMD in the null cells compared to controls (Fig. 1I, Supplemental Fig. 1J-K). Importantly, NLRP3 levels were similarly expressed in both genotypes, suggesting that increased caspase-1 activation was not a reflection of increased NLRP3 levels (Fig. 1F, Supplemental Fig. 1H).

### Tmem178 controls NLRP3 inflammasome activation via SOCE

We previously reported that Tmem178 deficiency leads to increased intracellular Ca^2+^ levels in myeloid cells (18). To determine whether Tmem178 modulates NLRP3 inflammasome via a calcium-dependent pathway, we cultured WT and *Tmem178*^-/-^ BMDMs in 2mM Ca^2+^, Ca^2+^ free media or with the calcium chelator BAPTA. As expected, the percent of cells with active caspase-1 and IL-1β release were higher in *Tmem178*^*-/-*^ cultures in the presence of physiological calcium levels (Supplemental Fig. 2A-D). However, both WT and *Tmem178*^*-/-*^ cells had barely detectable active caspase-1 and IL-1β release in Ca^2+^ free media or in the presence of BAPTA. While this result confirms that Ca^2+^ plays a key role in NLRP3 inflammasome signaling, it does not explain how Tmem178 deficiency regulates this pathway.

Since Tmem178 binds to Stim1 (18), a critical component of SOCE and modulator of intracellular Ca^2+^ levels (26) (27), next we wondered whether Tmem178 modulates inflammasome activation through SOCE. shRNA decreased Stim1 levels in WT and *Tmem178*^*-/-*^ BMDMs, but did not affect pro-IL-1β and pro-caspase-1 protein and transcript levels (Fig. 2A, Supplemental Fig. 3A-B), as expected. Interestingly, caspase-1 positive cells and IL-1β release were significantly reduced in Stim1-knockdown *Tmem178*^*-/-*^ but not WT BMDMs (Fig. 2B-D), indicating that SOCE drives NLRP3 inflammation activation in the context of *Tmem178* deficiency.

**Figure 2.**
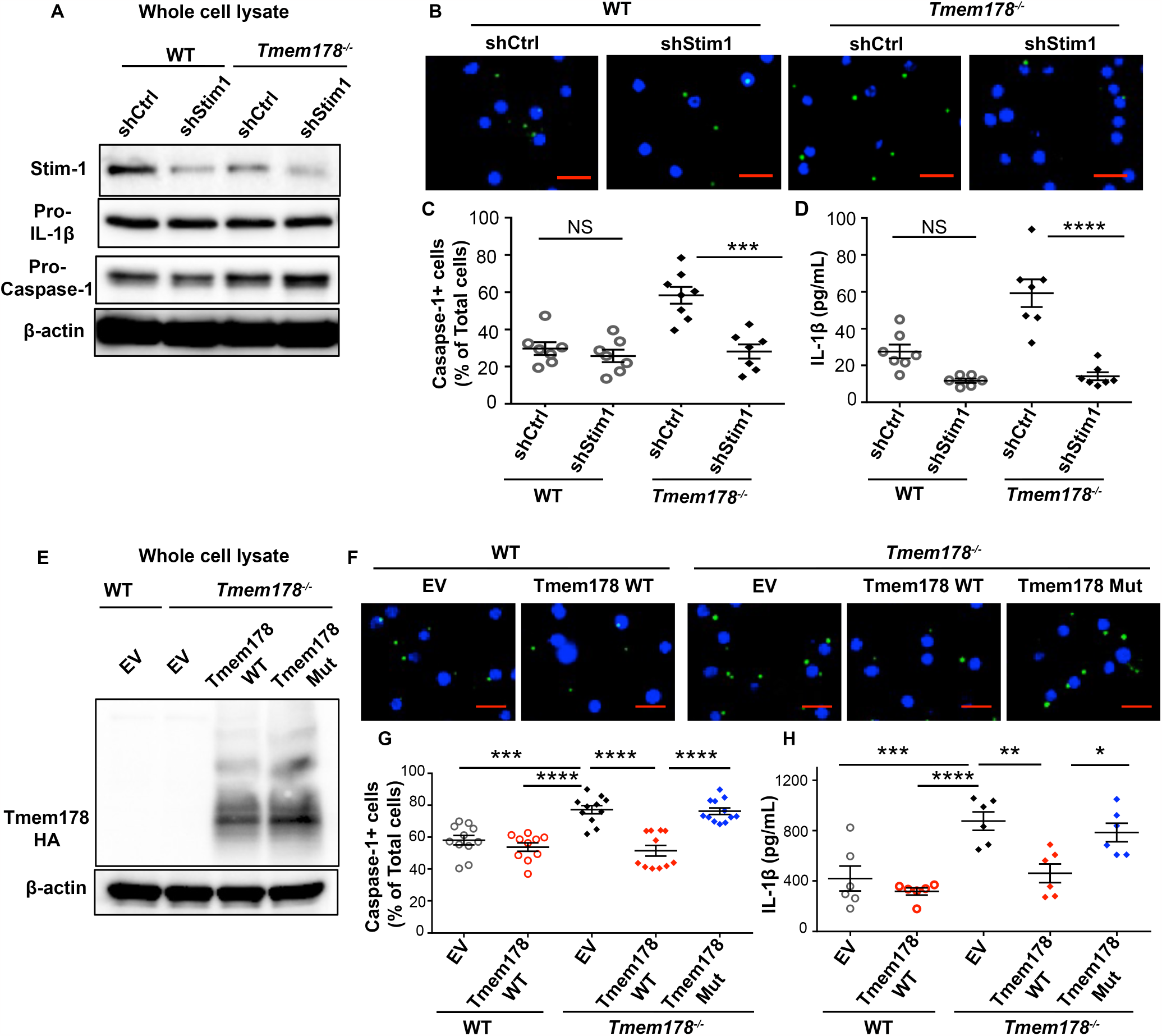
Tmem178 controls NLRP3 inflammasome activation via SOCE. WT and *Tmem178*^*-/-*^ BMDMs infected with shRNA-control or shRNA-Stim1 were (A) subjected to immunoblotting to measure Stim1, pro-IL-1β and pro-caspase-1. (B-C) Representative images of cells with active caspase-1 (green) following LPS (100ng/ml, 4hr) and nigericin (15μM, 45min) treatments (B). Quantification of percentage of cells with active caspase-1 (n = 7-8/group) (C) and IL-1β levels in culture supernatants by ELISA (n = 6-7/group) (D). (E-H) WT or *Tmem178*^*-/-*^ BMDMs infected with empty vector (EV), HA-tagged full length Tmem178 (Tmem178-WT) or mutant lacking Stim1 binding site (Tmem178-Mut) were (E) subjected to immunoblotting using anti-HA antibody. (F-H) Representative images of cells with active caspase-1 (green) following 100ng/ml LPS for 4hr, and 15μM nigericin for 45min (Scale=10μm) (F). Quantification of percentage of cells with active caspase-1 (n = 10-12) (G). IL-1β levels in culture supernatants measured by ELISA (n = 6) (H). Data are presented as mean ± SEM. Nuclei were stained with DAPI (blue). Statistical significance was determined by two-way ANOVA for multiple comparisons. \**P* < 0.05, \*\**P* < 0.01, \*\*\**P* < 0.001 and \*\*\*\**P* < 0.0001.

To better understand how Tmem178 controls NLRP3 inflammasome activation via Stim1, we expressed HA-tagged Tmem178-WT or the Tmem178 mutant lacking the Stim1 binding site (Tmem178-L212W;M216W as described in (18)) in *Tmem178*^*-/-*^ BMDMs. First, we confirmed similar protein and mRNA levels of *Tmem178* and *caspase-1* in the infected cells (Fig. 2E, Supplemental Fig. 3C-D). Next, we observed that expression of Tmem178-WT did not alter inflammasome activation in WT cells but reduced the percentage of caspase-1+ cells and IL-1β release in *Tmem178*^*-/-*^ BMDMs (Fig. 2F-H). Importantly, the expression of Tmem178 mutant lacking Stim1 binding in *Tmem178*^*-/-*^ cells restored active caspase-1 and IL-1β release to empty vector levels (Fig. 2F-H). These results indicate that loss of a direct interaction between Stim1 and Tmem178 is required to increase NLRP3 activation.

### Loss of Tmem178 induces mitochondria dysfunction

Mitochondrial damage is a known intracellular activator of inflammasomes, triggered by calcium overload (28). Because *Tmem178*^*-/-*^ BMDMs have increased basal intracellular calcium (Fig 3A-B and (18)), we wondered whether treatment with nigericin could further increase calcium levels following SOCE activation. WT and *Tmem178*^*-/-*^ BMDMs plated in calcium free medium with or without nigericin were stimulated with 2mM calcium to activate SOCE. While nigericin per se did not change calcium levels, SOCE activation led to a much higher calcium increase, as measured by the area under the curve (AUC), in nigericin-treated *Tmem178*^*-/-*^ BMDMs compared with WT (Fig. 3C-D).

**Figure 3.**
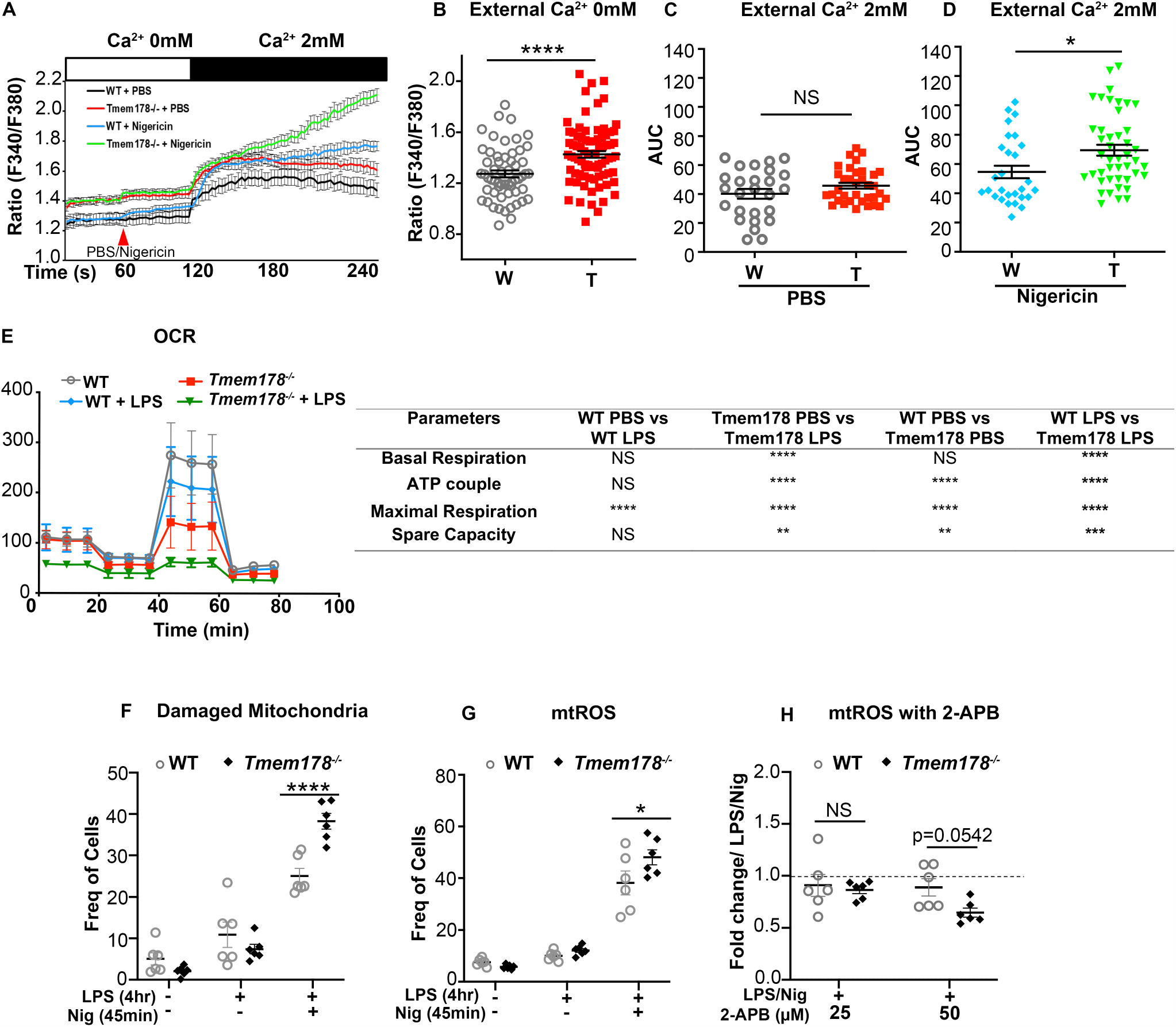
Loss of Tmem178 induces mitochondria dysfunction. (A-D) Calcium traces and area under curve (AUC) in WT and *Tmem178*^*-/-*^ BMDMs in calcium free medium stimulated with nigericin or PBS followed by 2mM calcium. (E) Measurement of oxygen consumption rate (OCR) before and after injections of oligomycin, FCCP and antimycin A in resting and LPS-stimulated (100ng/ml, 4hr) WT and *Tmem178*^*-/-*^ BMDMs. (F-G) Quantification of WT and *Tmem178*^*-/-*^ BMDMs following LPS (100ng/ml, 4hr) and nigericin (15μM, 45 min) for (F) damaged mitochondria (MitoGreen^high^ and MitoRed^low^) (n = 6/group) and (G) mtROS (MitoSOX^high^) (n = 6/group). (H) Fold change values of MitoSOX^high^ WT and *Tmem178-/-* BMDMs treated with 2-APB (25μM and 50μM for 30 min) after priming with LPS (100ng/ml, 4 hr) and nigericin (15μM, 45 min) (n = 6/group). MitoSOX^high^ cells exposed to LPS and nigericin were used as basal controls. Data are presented as mean ± SEM. Statistical significance was determined by two-tailed t-test (B, C, D) or two-way ANOVA for multiple comparisons (E-H). \**P* < 0.05 and \*\*\*\**P* < 0.0001.

To examine the effects of Tmem178 loss on mitochondria fitness, we measured the Oxygen Consumption Rate (OCR). Strikingly, *Tmem178*^*-/-*^ BMDMs had reduced OCR in basal conditions and after LPS simulation compared with WT (Fig. 3E). Next, we stained BMDMs with MitoTracker Red that is taken up by negatively charged mitochondria and, thus is used to determine mitochondrial functionality (29). We also used MitoTracker Green, which reacts with cysteine residues on mitochondrial proteins and is used to represent mitochondrial mass (30). At steady state, we did not detect differences between WT and *Tmem178*^*-/-*^ BMDMs, with less than 6% of cells having damaged mitochondria (Fig. 3F). LPS increased mitochondria damage in both genotypes, but it was further enhanced in *Tmem178*^*-/-*^ BMDMs after nigericin stimulation (Fig. 3F, Supplemental Fig. 4A-B). Next, we measured mtROS, a known activator of the NLRP3 inflammasome (31), by staining the cells with the MitoSOX dye. Consistent with elevated mitochondria damage, mtROS levels were higher in *Tmem178*^*-/-*^ cells treated with LPS and nigericin compared with WT (Fig. 3G, Supplemental Fig. 4C-D).

Finally, to determine if increased SOCE activation induces mitochondria dysfunction in *Tmem178*^*-/-*^ cells, we treated BMDMs with different doses of 2-Aminoethoxydiphenyl borate (2-APB) to inhibit SOCE, and measured mtROS levels. Strikingly, 2-APB significantly reduced mtROS in *Tmem178*^*-/-*^ cells but not WT (Fig. 3H). These findings indicate that loss of Tmem178 activates SOCE and induces mitochondrial damage and mtROS, events that lead to NLRP3 inflammasome activation.

### Inhibition of inflammasome signaling ameliorates CSS in *Tmem178*^*-/-*^ mice

To further demonstrate the importance of Tmem178 in modulating IL-1β production via the NLRP3 inflammasome *in vivo*, we generated WT, *Tmem178*^*-/-*^, *Nlrp3*^*-/*^, *Tmem178*^*-/-*^;*Nlrp3*^*-/-*^ mice and examined IL-1β circulating levels following 6-hour stimulation with 10mg/kg LPS. Confirming our initial observations, IL-1β levels in the serum were significantly increased in *Tmem178*^*-/-*^ mice compared to WT, while barely detectable in *Nlrp3*^*-/-*^ and *Tmem178*^*-/-*^;*/Nlrp3*^*-/-*^ mice (Fig. 4A). These results indicate that loss of Tmem178 requires the NLRP3 inflammasome for IL-1β secretion, thus excluding the involvement of other inflammasomes.

**Figure 4.**
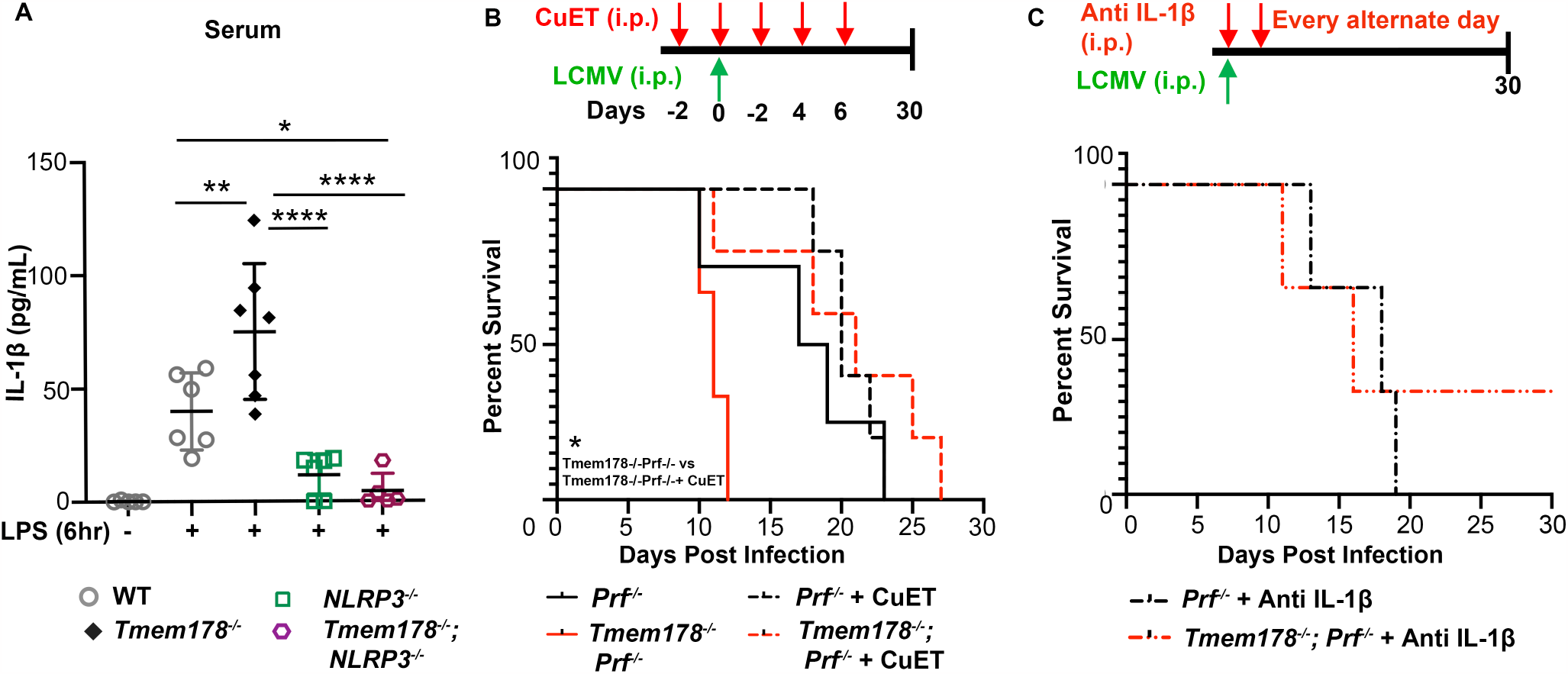
Inhibition of inflammasome signaling ameliorates CSS in *Tmem178*^*-/-*^ mice. (A) IL-1β levels in serum measured by ELISA in WT, *Tmem178*^*-/-*^, *Nlrp3*^*-/-*^, and *Tmem178*^*-/-*^*;Nlrp3*^*-/-*^ mice following i.p. injection with 10mg/kg LPS or PBS for 6hr prior to sample collection (n = 5-7). (B-C) Percent of survival of LCMV-infected *Prf*^*-/-*^ mice or *Tmem178*^*-/-*^*;Prf*^*-/-*^ treated with (B) CuET (1mg/Kg i.p.; n = 5/group), starting 2 days prior to LCMV infection and continued every other day for a total of 5 doses. Mice receiving vehicle were used as control (n = 3 *Tmem178*^*-/-*^*;Prf*^*-/-*^ and n = 4 *Prf*^*-/-*^ mice) or (C) anti-IL-1β (10μg, n = 3/group) starting 4hr after LCMV-infection and continued every other day for 30 days. Data are presented as mean ± SEM. Statistical significance was determined by two-way ANOVA (A), Log-rank (Mantel-Cox) test (B, C). \**P* < 0.05, \*\**P* < 0.01 and \*\*\*\**P* < 0.0001.

Next, we used a well-established CSS mouse model consisting of *Perforin (Prf)*^-/-^ mice infected with LCMV, which develop severe macrophage-driven inflammatory responses due to the inability of T cells to clear the virus (32). To inhibit inflammasome activation, mice were administered bis(diethyldithiocarbamate)-copper (CuET), the active metabolite of disulfiram, a compound in current use to inhibit GSDMD, the common effector of inflammasomes, including the NLRP3 inflammasome (33). Animals were administered with 1mg/Kg CuET or vehicle given every alternate day, starting 2 days prior up to 6 days post virus infection. Similar to our previous finding, *Prf*^*-/-*^*;Tmem178*^*-/-*^ mice were highly susceptible to LCMV infection, with all animals succumbing within 10-13 days, as compared to *Prf*^*-/-*^ which died within 20-22 days. CuET did not improve the overall survival of LCMV-infected *Prf*^*-/-*^ animals, while significantly prolonged the life of *Prf*^*-/-*^*;Tmem178*^*-/-*^ animals for additional 2 weeks (Fig. 4B). Confirming that CuET can target GSDMD downstream of the NLRP3 inflammasome, IL-1β levels were significantly reduced by CuET in *Tmem178*^*-/-*^ BMDM following stimulation with LPS and nigericin (Supplemental Fig. 5). Furthermore, neutralizing IL-1 antibody, administered every other day for the duration of the experiment, also increased the survival of *Prf*^*-/-*^*;Tmem178*^*-/-*^ similar to *Prf*^*-/-*^ mice (Fig. 4C). Collectively, these results support the contribution of the NLRP3 inflammasome in the development of CSS in the context of *Tmem17*8 deficiency.

### Loss of Tmem178 regulates IL-1β release in sJIA

sJIA patients have elevated levels of IL-1β in circulation. To investigate whether increased IL-1B expression correlates with reduced *TMEM178* levels, we analyzed a published data set of human monocytes from sJIA patients and healthy donors ((34), GSE147608). We confirmed significant expression of *IL1B, STIM1* and *CASPASE-1* and a significant reduction of *TMEM178A* in the patients compared to healthy donors (Fig 5A-B). Similarly, the THP1 human monocyte cell line exposed to plasma from sJIA patients showed reduced *TMEM178* levels (Fig. 5C). Regression analysis further showed an inverse relation between *Tmem178* mRNA and IL-1β levels in WT BMDMs treated with sJIA plasma from different patients versus healthy controls. Further demonstrating that reduced *Tmem178* expression drives IL-1β release, WT and *Tmem178*^*-/-*^ BMDMs were incubated with healthy or sJIA plasma and stimulated with LPS and nigericin. IL-1β release in WT cells was increased by sJIA plasma (Fig 5D), a condition that reduces *Tmem178* expression (16). Interestingly, IL-1β levels were equally elevated by healthy or sJIA plasma in *Tmem178*^*-/-*^ BMDMs (Fig. 5E). These findings indicate that downregulation of Tmem178 is needed to induce IL-1β release and this could be the leading cause of elevated IL-1β levels during sJIA pathogenesis.

**Figure 5.**
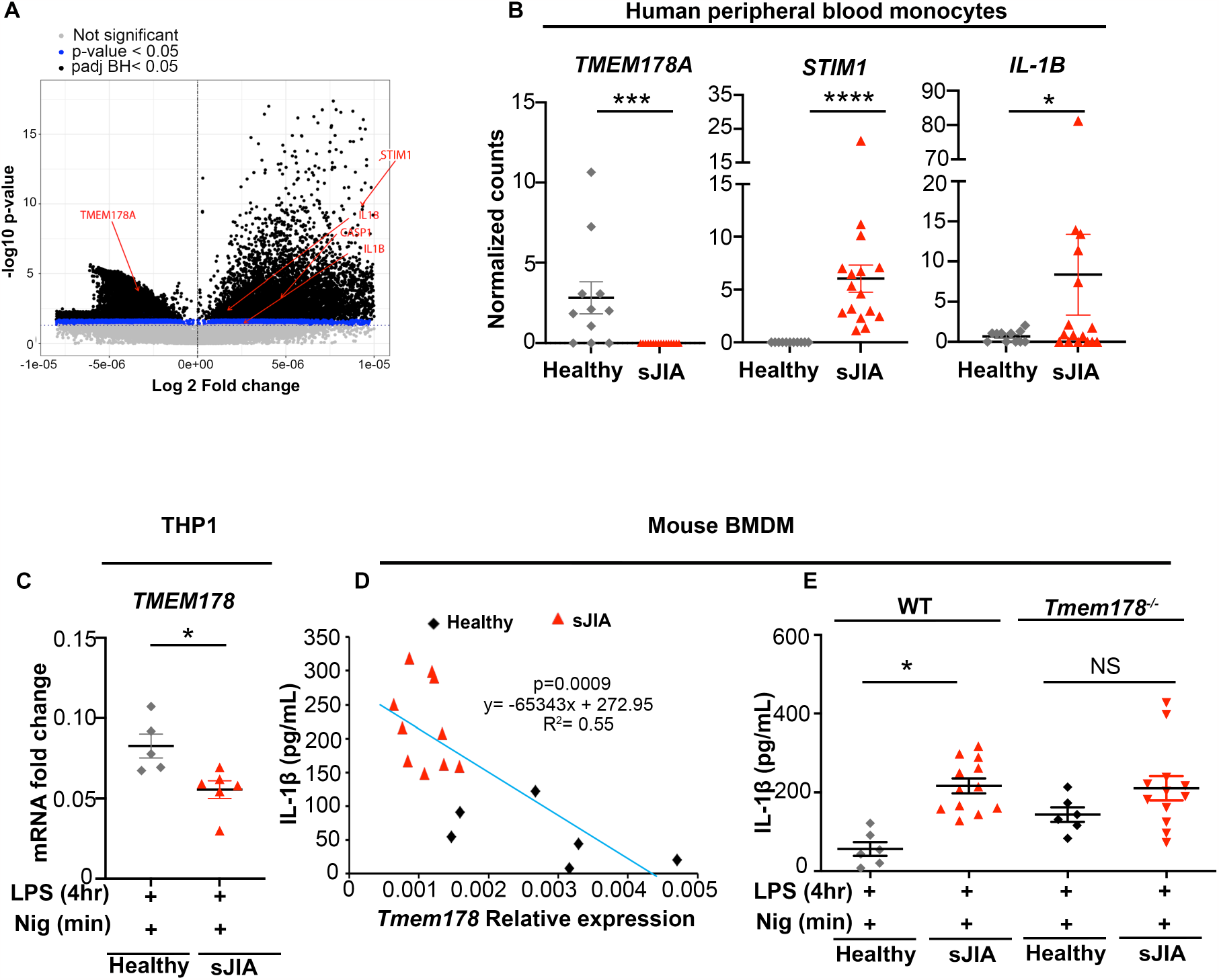
Tmem178 downregulation modulates IL-1β release in sJIA. (A-B) Differential gene expression in human peripheral blood monocytes from sJIA patients (n = 16) versus healthy controls (n = 11). (A) Volcano plot with genes of interest shown in red; P-value_adjusted(BH)_< 0.05 shown in black; P-value_unadjusted_< 0.05 in blue; genes with no significant difference in grey. (B) Normalized gene counts. (C) *TMEM178* mRNA in THP1 cells exposed to sJIA or healthy plasma (16 hr), followed by LPS (100ng/ml, 4 hr) and nigericin (7.5μM, 45min). (D) Regression analysis showing *Tmem178* mRNA relative to IL-1β by ELISA in WT BMDMs treated with sJIA versus healthy plasma (16 hr), followed by LPS (100ng/ml, 4 hr) and nigericin (7.5μM, 45min). (E) IL-1β ELISA in WT and *Tmem178*^*-/-*^ BMDM treated with healthy (n = 6) or sJIA (n = 12) plasma (16 hr), followed by LPS (100ng/ml, 4hr) and nigericin (7.5μM, 45min). Data are presented as mean ± SEM. Statistical significance was determined by Wald test using DESeq2 (A-B), two-tailed *t*-test (C) or two-way ANOVA for multiple comparisons (E). \**P* < 0.05, \*\*\**P* < 0.001, \*\*\*\**P* < 0.0001.

## Discussion

Identifying sJIA patients who might develop severe inflammation associated with cytokine storm syndrome and understanding the etiology of CSS are unmet clinical needs. Various diagnostic tools have been developed for identifying CSS in the setting of sJIA based on clinical (e.g., fever) and laboratory features (e.g., cytopenia) (35). However, none is reliably sensitive and specific. Here, we have identified Tmem178 as a negative regulator of inflammasome activation, whose expression levels in monocytes/macrophages negatively correlate with IL-1β in sJIA. Most importantly, we report that the premature death of *Tmem178*^*-/-*^ mice in a LCMV-induced CSS model can be prevented by inflammasome inhibition. Our data supports *Tmem178* expression levels as a biomarker of sJIA/CSS progression and show the efficacy of targeting of inflammasomes in the context of reduced Tmem178 in macrophages.

Elevated IL-1β and IL-18 have been observed in sJIA and CSS (36). While blocking IL-1β alone by canakinumab does not reduce the risk to develop CSS in sJIA patients; anakinra, a recombinant IL-1 receptor antagonist, induces remission with normalization of lab abnormalities and fever to some patients poorly responsive to more traditional therapies (37) (38). Free IL-18 is also highly elevated in MAS patients. Blockade of IL-18 receptor reduces inflammation in a murine model of CSS (8). Importantly IL-18 inhibition in combination with anakinra successfully improved life-threatening hyperinflammation in a patient with a dominant heterozygous mutation in the NLRC4 inflammasome (13). These clinical funding suggests that aberrant activation of inflammasomes could be driving disease progression in sJIA patients developing CSS.

The causes of elevated IL-1β and IL18 levels in sJIA and CSS are not known. A gain-of-function mutation of *NLRC4* (13), and a heterozygous missense variant in the *CASP1* gene encoding pro-caspase-1 have been reported to induce severe recurrent CSS in patients with sJIA (12). However, activating-mutations in inflammasomes or their components are very rare. Upregulation of AIM2 and NLRC4 inflammasomes have been detected in neutrophils from children with both active and inactive sJIA (39). Elevation in S100 proteins, known for priming the NLRP3 inflammasome, has also been reported further supporting inflammasome involvement in sJIA (39).

In this study, we show that downregulation of *Tmem178* levels drives IL-1β production via SOCE-dependent NLRP3 inflammasome activation in macrophages. We find increased IL-1β transcripts in *Tmem178*^*-/-*^ cells, although NF-kB activation, *NLRP3* levels, and cytoplasmic pro-IL-1β levels are similar to WT. This apparent controversy could be dependent on a more efficient cleavage of pro-IL-1β by caspase-1. The increased caspase-1 positive cells in *Tmem178*^*-/-*^ cultures and higher levels of GSDMD, the pore forming protein involved in IL-1β secretion, strongly support this hypothesis.

Our findings are in line with a previous gene profiling study in PBMCs from patients with sJIA with early-stage CSS that identified a cluster enriched for genes involved in innate immune responses (40). Our results are clinically relevant as we find reduced *TMEM178* while increased *IL-1B* levels in a human data set of monocytes from sJIA patients. Our finding from naïve human and murine macrophages exposed to sJIA plasma also shows similar results. The release of IL-1β in BMDMs exposed to sJIA plasma is dependent on downregulation of *Tmem178*. Confirming the hypothesis that downregulation of *Tmem178* drives NLRP3 activation and possibly increases the susceptibility to CSS, we find that IL-1β neutralization or inflammasome inhibition significantly increase the survival of *Prf*^*-/-*^*;Tmem178*^*-/-*^ mice to LCMV infection. Considering that loss of Tmem178 in macrophages upregulates several inflammatory cytokines (16), the prolonged survival to CSS confirms a key role for IL-1β and possibly IL-18 in disease severity in the context of *Tmem178* deficiency.

Numerous studies have shown that calcium regulates NLRP3 inflammasome activation by increasing the assembly of inflammasome components (41) (42). Both intracellular calcium release from the ER or mitochondria, and extracellular calcium entry via GPCRs and calcium channels can activate the NLRP3 inflammasome (42), (43). These findings support our results using BAPTA, to chelate intracellular calcium, and calcium free medium, to remove the extracellular source of calcium. However, the role SOCE on NLRP3 inflammasome activation is controversial. While a recent publication indicated the involvement of the Stim1 SOCE component in inducing NLRP3 upregulation and cytokine production in lung epithelial cells infected with influenza virus (44), studies using *Stim1*^*-/-*^*/Stim2*^*-/-*^ mice indicated that SOCE is dispensable for NLRP3 inflammasome activation in macrophages (45). The novelty of our finding consists in demonstrating that increased SOCE activation in the context of Tmem178 deficiency drives NLRP3 inflammasome activation and IL-1β release in macrophages. Furthermore, we now provide new data linking increased calcium levels to mitochondria damage and release of mtROS, as the mechanism behind SOCE-mediated NLRP3 inflammasome activation.

Calcium overload has been shown to cause mitochondrial damage (28). We confirmed our previous observations that *Tmem178*^*-/-*^ BMDMs have higher basal calcium levels than WT. Interestingly, SOCE activation following nigericin leads to even higher increase in calcium in the null cells. It is very likely that elevated calcium in *Tmem178*^*-/-*^ BMDMs induces mitochondria damage, as shown by reduced OCR and increased mtROS. Consistent with this hypothesis, the SOCE inhibitor 2-APB reduces mtROS levels in *Tmem178*^*-/-*^ but not WT BMDMs. Mitochondrial damage and mtROS have been implicated in NLRP3 activation (31). Recently, increased cleaved GSDMD and its insertion into the mitochondria has been also related to mtROS generation and IL-1β maturation in Thp1 macrophages (46). Similarly, we find increased GSDMD cleavage in *Tmem178*^*-/-*^ cells. Furthermore, loss of GSDMD confers protective effects in a mouse model of CSS (47). Altogether, our data provide new mechanistic insights into NLRP3 activation via downregulation of Tmem178, increased SOCE and cleavage of GSDMD, and induction of mtROS (Fig. 6).

**Figure 6.**
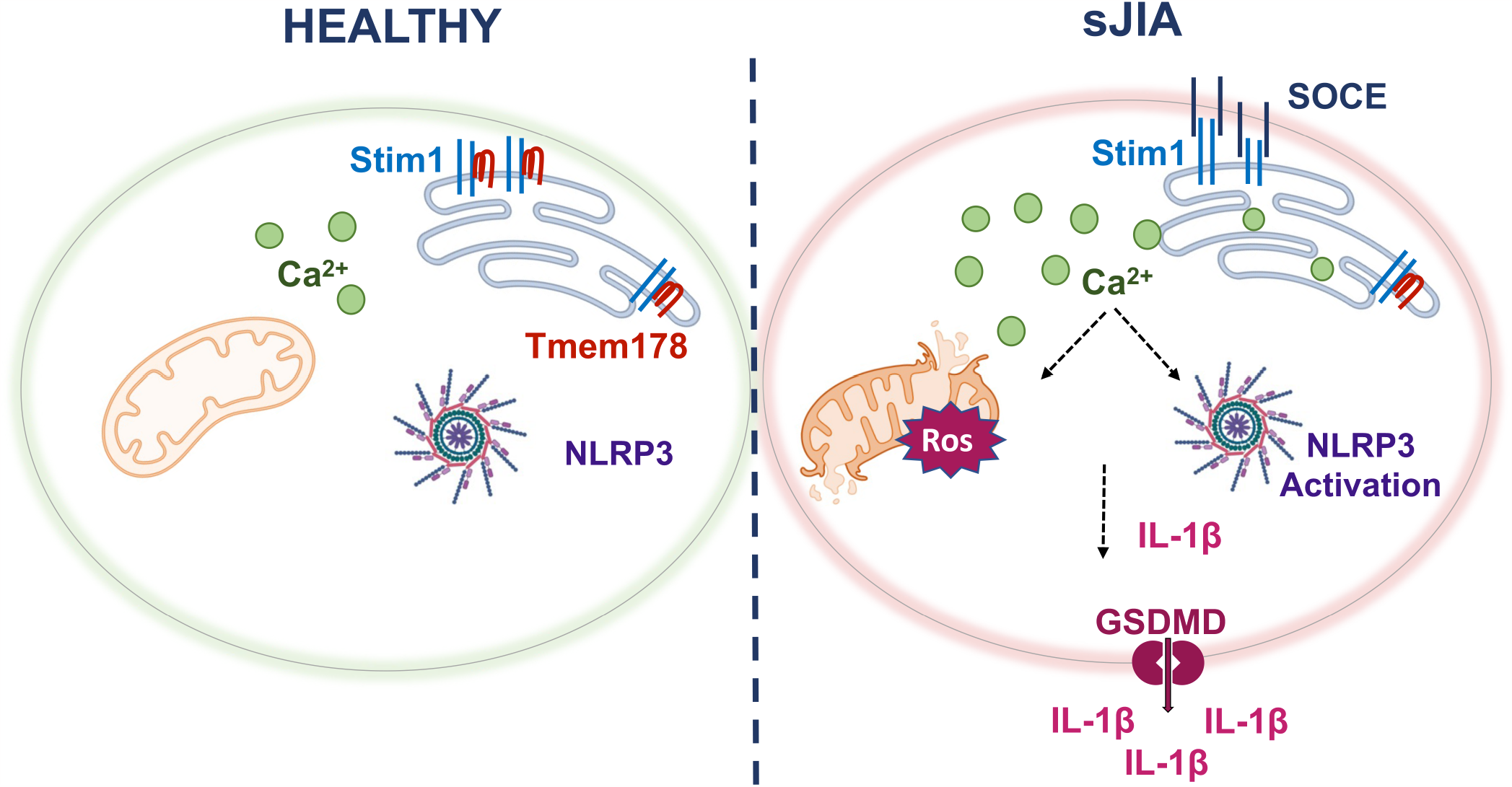
Model of Tmem178-mediated regulation of IL-1β levels in sJIA. Downregulation of Tmem178 in sJIA drives the activation of SOCE and increases intracellular calcium levels. Calcium overload leads to mitochondria damage and the release of mtROS, that ultimately activate NLRP3 inflammasome assembly. Increased *pro-IL-1β* production and generation of mature IL-1β by caspase-1 cleavage, together with its secretion through the pore forming protein GSDMD, leads to excessive release of IL-1β in circulation in sJIA pathologies.

In the recent era of COVID-19, numerous research studies showing disease severity in COVID-19 patients suggests the involvement of NLRP3 inflammasome, activation of key inflammatory molecules like active caspase-1, and elevated levels of IL-18 in the sera/tissue or PBMCs samples (14). Considering that loss of *Tmem178* can drive exuberant NLRP3 inflammasome activation, it would be important to monitor *Tmem178* levels in COVID-19 patients to determine the risk of developing CSS. In conclusion, herein we report a novel mechanism of NLRP3 inflammasome/IL-1β regulation via loss of Tmem178 through modulation of SOCE and mitochondria damage, suggesting that *Tmem178* levels could be used as a biomarker of disease severity and inflammasomes could be considered a potential therapeutic target for sJIA, CSS or other inflammatory disorders accompanied by loss of *Tmem178* expression in macrophages.

## Acknowledgements

This work was supported by the Shriners Hospital Grant 85170 (to R.F.), NIH Grants R01 AR066551 and CA235096 (to R.F.), and Siteman Cancer Center (to R.F.), NIH/NIAMS AR076758 and AI161022 grants (to G.M.) and by the NIH P30 Grants AR057235 and P30 AR074992. We thank Dr. Elizabeth Mellins from Stanford University for providing patient plasma samples.

## References

1. Grom AA, Horne A, De Benedetti F. Macrophage activation syndrome in the era of biologic therapy. Nat Rev Rheumatol. 2016;12(5):259–68.

2. Yasin S, Schulert GS. Systemic juvenile idiopathic arthritis and macrophage activation syndrome: update on pathogenesis and treatment. Curr Opin Rheumatol. 2018;30(5):514–20.

3. Bracaglia C, Prencipe G, De Benedetti F. Macrophage Activation Syndrome: different mechanisms leading to a one clinical syndrome. Pediatr Rheumatol Online J. 2017;15(1):5.

4. Merad M, Martin JC. Pathological inflammation in patients with COVID-19: a key role for monocytes and macrophages. Nat Rev Immunol. 2020;20(6):355–62.

5. Shimabukuro-Vornhagen A, Gödel P, Subklewe M, Stemmler HJ, Schlößer HA, Schlaak M, et al. Cytokine release syndrome. J Immunother Cancer. 2018;6(1):56.

6. Sharma C, Ganigara M, Galeotti C, Burns J, Berganza FM, Hayes DA, et al. Multisystem inflammatory syndrome in children and Kawasaki disease: a critical comparison. Nat Rev Rheumatol. 2021;17(12):731–48.

7. Toplak N, Blazina Š, Avčin T. The role of IL-1 inhibition in systemic juvenile idiopathic arthritis: current status and future perspectives. Drug Des Devel Ther. 2018;12:1633–43.

8. Girard-Guyonvarc’h C, Palomo J, Martin P, Rodriguez E, Troccaz S, Palmer G, et al. Unopposed IL-18 signaling leads to severe TLR9-induced macrophage activation syndrome in mice. Blood. 2018;131(13):1430–41.

9. Weiss ES, Girard-Guyonvarc’h C, Holzinger D, de Jesus AA, Tariq Z, Picarsic J, et al. Interleukin-18 diagnostically distinguishes and pathogenically promotes human and murine macrophage activation syndrome. Blood. 2018;131(13):1442–55.

10. Broz P, Dixit VM. Inflammasomes: mechanism of assembly, regulation and signalling. Nat Rev Immunol. 2016;16(7):407–20.

11. Man SM, Kanneganti TD. Converging roles of caspases in inflammasome activation, cell death and innate immunity. Nat Rev Immunol. 2016;16(1):7–21.

12. Jørgensen SE, Christiansen M, Høst C, Glerup M, Mahler B, Lausten MM, et al. Systemic juvenile idiopathic arthritis and recurrent macrophage activation syndrome due to a CASP1 variant causing inflammasome hyperactivation. Rheumatology (Oxford). 2020;59(10):3099–105.

13. Canna SW, de Jesus AA, Gouni S, Brooks SR, Marrero B, Liu Y, et al. An activating NLRC4 inflammasome mutation causes autoinflammation with recurrent macrophage activation syndrome. Nat Genet. 2014;46(10):1140–6.

14. Vora SM, Lieberman J, Wu H. Inflammasome activation at the crux of severe COVID-19. Nat Rev Immunol. 2021;21(11):694–703.

15. Decker CE, Yang Z, Rimer R, Park-Min KH, Macaubas C, Mellins ED, et al. Tmem178 acts in a novel negative feedback loop targeting NFATc1 to regulate bone mass. Proc Natl Acad Sci U S A. 2015;112(51):15654–9.

16. Mahajan S, Decker CE, Yang Z, Veis D, Mellins ED, Faccio R. Plcγ2/Tmem178 dependent pathway in myeloid cells modulates the pathogenesis of cytokine storm syndrome. J Autoimmun. 2019;100:62–74.

17. Takeshita S, Kaji K, Kudo A. Identification and characterization of the new osteoclast progenitor with macrophage phenotypes being able to differentiate into mature osteoclasts. J Bone Miner Res. 2000;15(8):1477–88.

18. Yang Z, Yan H, Dai W, Jing J, Yang Y, Mahajan S, et al. Tmem178 negatively regulates store-operated calcium entry in myeloid cells via association with STIM1. J Autoimmun. 2019;101:94–108.

19. Benjamini YaH, Y., Journal of the Royal statistical society: series B (Methodological), 57 (1), pp.289–300. Controlling the false discovery rate: a practical and powerful approach to multiple testing.: Journal of the Royal statistical society: series B (Methodological); 1995. p. pp.289-300.

20. Karin M, Ben-Neriah Y. Phosphorylation meets ubiquitination: the control of NF-[kappa]B activity. Annu Rev Immunol. 2000;18:621–63.

21. Yang CA, Huang ST, Chiang BL. Association of NLRP3 and CARD8 genetic polymorphisms with juvenile idiopathic arthritis in a Taiwanese population. Scand J Rheumatol. 2014;43(2):146–52.

22. Ratajczak MZ, Kucia M. SARS-CoV-2 infection and overactivation of Nlrp3 inflammasome as a trigger of cytokine “storm” and risk factor for damage of hematopoietic stem cells. Leukemia. 2020;34(7):1726–9.

23. Galliher-Beckley AJ, Lan LQ, Aono S, Wang L, Shi J. Caspase-1 activation and mature interleukin-1β release are uncoupled events in monocytes. World J Biol Chem. 2013;4(2):30–4.

24. Gurung P, Li B, Subbarao Malireddi RK, Lamkanfi M, Geiger TL, Kanneganti TD. Chronic TLR Stimulation Controls NLRP3 Inflammasome Activation through IL-10 Mediated Regulation of NLRP3 Expression and Caspase-8 Activation. Sci Rep. 2015;5:14488.

25. den Hartigh AB, Fink SL. Detection of Inflammasome Activation and Pyroptotic Cell Death in Murine Bone Marrow-derived Macrophages. J Vis Exp. 2018(135).

26. Zhang SL, Yu Y, Roos J, Kozak JA, Deerinck TJ, Ellisman MH, et al. STIM1 is a Ca2+ sensor that activates CRAC channels and migrates from the Ca2+ store to the plasma membrane. Nature. 2005;437(7060):902–5.

27. Roos J, DiGregorio PJ, Yeromin AV, Ohlsen K, Lioudyno M, Zhang S, et al. STIM1, an essential and conserved component of store-operated Ca2+ channel function. J Cell Biol. 2005;169(3):435–45.

28. Brookes PS, Yoon Y, Robotham JL, Anders MW, Sheu SS. Calcium, ATP, and ROS: a mitochondrial love-hate triangle. Am J Physiol Cell Physiol. 2004;287(4):C817–33.

29. Buckman JF, Hernández H, Kress GJ, Votyakova TV, Pal S, Reynolds IJ. MitoTracker labeling in primary neuronal and astrocytic cultures: influence of mitochondrial membrane potential and oxidants. J Neurosci Methods. 2001;104(2):165–76.

30. Xiao B, Deng X, Zhou W, Tan EK. Flow Cytometry-Based Assessment of Mitophagy Using MitoTracker. Front Cell Neurosci. 2016;10:76.

31. Zhou R, Yazdi AS, Menu P, Tschopp J. A role for mitochondria in NLRP3 inflammasome activation. Nature. 2011;469(7329):221–5.

32. Jordan MB, Hildeman D, Kappler J, Marrack P. An animal model of hemophagocytic lymphohistiocytosis (HLH): CD8+ T cells and interferon gamma are essential for the disorder. Blood. 2004;104(3):735–43.

33. Wang C, Yang T, Xiao J, Xu C, Alippe Y, Sun K, et al. NLRP3 inflammasome activation triggers gasdermin D-independent inflammation. Sci Immunol. 2021;6(64):eabj3859.

34. Schulert GS, Pickering AV, Do T, Dhakal S, Fall N, Schnell D, et al. Monocyte and bone marrow macrophage transcriptional phenotypes in systemic juvenile idiopathic arthritis reveal TRIM8 as a mediator of IFN-γ hyper-responsiveness and risk for macrophage activation syndrome. Ann Rheum Dis. 2021;80(5):617–25.

35. Boom V, Anton J, Lahdenne P, Quartier P, Ravelli A, Wulffraat NM, et al. Evidence-based diagnosis and treatment of macrophage activation syndrome in systemic juvenile idiopathic arthritis. Pediatr Rheumatol Online J. 2015;13:55.

36. Shimizu M. Macrophage activation syndrome in systemic juvenile idiopathic arthritis. Immunol Med. 2021;44(4):237–45.

37. Sönmez HE, Demir S, Bilginer Y, Özen S. Anakinra treatment in macrophage activation syndrome: a single center experience and systemic review of literature. Clin Rheumatol. 2018;37(12):3329–35.

38. Rajasekaran S, Kruse K, Kovey K, Davis AT, Hassan NE, Ndika AN, et al. Therapeutic role of anakinra, an interleukin-1 receptor antagonist, in the management of secondary hemophagocytic lymphohistiocytosis/sepsis/multiple organ dysfunction/macrophage activating syndrome in critically ill children*. Pediatr Crit Care Med. 2014;15(5):401–8.

39. Brown RA, Henderlight M, Do T, Yasin S, Grom AA, DeLay M, et al. Neutrophils From Children With Systemic Juvenile Idiopathic Arthritis Exhibit Persistent Proinflammatory Activation Despite Long-Standing Clinically Inactive Disease. Front Immunol. 2018;9:2995.

40. Fall N, Barnes M, Thornton S, Luyrink L, Olson J, Ilowite NT, et al. Gene expression profiling of peripheral blood from patients with untreated new-onset systemic juvenile idiopathic arthritis reveals molecular heterogeneity that may predict macrophage activation syndrome. Arthritis Rheum. 2007;56(11):3793–804.

41. Lee GS, Subramanian N, Kim AI, Aksentijevich I, Goldbach-Mansky R, Sacks DB, et al. The calcium-sensing receptor regulates the NLRP3 inflammasome through Ca2+ and cAMP. Nature. 2012;492(7427):123–7.

42. Murakami T, Ockinger J, Yu J, Byles V, McColl A, Hofer AM, et al. Critical role for calcium mobilization in activation of the NLRP3 inflammasome. Proc Natl Acad Sci U S A. 2012;109(28):11282–7.

43. Rossol M, Pierer M, Raulien N, Quandt D, Meusch U, Rothe K, et al. Extracellular Ca2+ is a danger signal activating the NLRP3 inflammasome through G protein-coupled calcium sensing receptors. Nat Commun. 2012;3:1329.

44. Liu CC, Miao Y, Chen RL, Zhang YQ, Wu H, Yang SM, et al. STIM1 mediates IAV-induced inflammation of lung epithelial cells by regulating NLRP3 and inflammasome activation via targeting miR-223. Life Sci. 2021;266:118845.

45. Vaeth M, Zee I, Concepcion AR, Maus M, Shaw P, Portal-Celhay C, et al. Ca2+ Signaling but Not Store-Operated Ca2+ Entry Is Required for the Function of Macrophages and Dendritic Cells. J Immunol. 2015;195(3):1202–17.

46. Platnich JM, Chung H, Lau A, Sandall CF, Bondzi-Simpson A, Chen HM, et al. Shiga Toxin/Lipopolysaccharide Activates Caspase-4 and Gasdermin D to Trigger Mitochondrial Reactive Oxygen Species Upstream of the NLRP3 Inflammasome. Cell Rep. 2018;25(6):1525-36.e7.

47. Tang S, Yang C, Li S, Ding Y, Zhu D, Ying S, et al. Genetic and pharmacological targeting of GSDMD ameliorates systemic inflammation in macrophage activation syndrome. J Autoimmun. 2022;133:102929.

