## Supplemental Information for "Tmem178 negatively regulates IL-1β production through inhibition of the NLRP3 inflammasome"

**Supplemental material and methods**

**Peritoneal Fluid Collection.** A needle was inserted through the skin of the abdomen into the peritoneal cavity of the anesthetized mice and a syringe containing 1mL of sterile PBS was attached to the end of the needle. The PBS was injected back and forth to collect the peritoneal fluid. Samples were stored at -80 $^0$C prior to use.

**Serum preparation and liver processing.** Blood was collected from the heart of the anesthetized mice and serum was separated using the 1.1ml Serum-Gel Polypropylene tubes (Thermo Fisher, USA) after 30 min incubation at room temperature. The tubes were centrifuged at 8,000 rpm for 4 min and the clear serum was collected and stored at -80 $^0$C until further use. The livers were immediately harvested without perfusion and processed as shown previously (16).

**Cell isolation and inflammasome activation.** To generate bone marrow-derived macrophages (BMDM), bone marrow cells flushed from femurs and tibias of 6-8 weeks old mice were cultured in α-minimum essential medium (α-MEM, Sigma, MO, USA) with 10% fetal bovine serum (GIBCO, NY, USA), 2mM glutamine (Corning, NY, USA), penicillin/streptomycin (100IU/ml) (GIBCO, NY, USA), and 10% CMG14-12 cell conditioned medium (17) as a source of M-CSF. This medium was used to maintain BMDMs in 100mm petri dishes for initial attachment and expansion of BMDMs. The cells were dissociated from the petri dishes using Trypsin-EDTA (0.25%) (Gibco, NY, USA). For certain experiments, BMDMs were starved overnight in serum and cytokine free medium, primed with 100ng/ml LPS (4 hours) followed by nigericin (15μM, 45 min) stimulation to activate the NLRP3 inflammasome. Typically, 0.5 million cells were cultured in 6-well plates (TPP, Switzerland) for RT-PCR or ELISA and 1 million cells were cultured in

60cm$^2$ tissue culture plates (TPP, Switzerland) for Immunoblotting. Culture supernatants were used for ELISA and cell lysates for RT-PCR and immunoblotting.

**Active caspase1 staining and imaging.** To measure active caspase-1, BMDMs were labeled with the FAM-YVAD-FMK probe using the FAM-FLICA caspase-1 assay kit (Immunochemistry Technologies). Briefly, BMDMs were cultured overnight at a density of 50,000 cells per well in a 96-well plate. Cells were then stimulated with 100ng/ml LPS for 4 hours and 15µM nigericin for 45 min followed by incubation with the FAM-YVAD-FMK probe for 30 min. Supernatants were collected to measure IL-1β release. Cells were washed, fixed with 10% buffered formalin and then counterstained with Fluoro-gel II containing DAPI (Electron Microscopy Sciences). Images were taken with a DP70 Olympus Digital Camera coupled to an IX51Olympus inverted microscope, captured with QCapture Pro software and analyzed with ImageJ software.

**Lentivirus and retrovirus preparation and cell transduction.** HEK293T cells were cultured in Dulbecco's modified Eagle's medium containing 10% heat-inactivated fetal bovine serum, 100 IU/ml penicillin plus 100µg/ml streptomycin, and 1mM sodium pyruvate (25- 000-Cl, Corning). shRNA Ctrl (targeting a scramble sequence) or shRNA Stim1 (targeting the CCCTTCCTTTCTTTGCAATAT sequence in the 3'-UTR region of *Stim1*) in pLKO.1 lentiviral vector, containing a puromycin resistance cassette, were co-transfected together with 1ng packaging plasmid Delta8.2 and 0.125µg envelop plasmid VSVg into HEK293T cells using the Polyjet transfection reagent (SL100688, SignaGen Laboratories). Supernatants containing the lentivirus were harvested 36-48 hours after transfection, filtered, and used to infect BMDMs. 50% lentivirus supernatant and 10% CMG 14-12 in α-10 medium with additional 8mg/ml Polybrene were added to the cells for 24 hours. Cells were then selected in α-10 medium containing 10%

CMG 14-12 as source of M-CSF and 2μg/ml puromycin for 24 hours prior to being used for the indicated experiments.

PLAT-E cells were plated in 6 cm dishes and transfected with 2μg of HA-tagged Tmem178 Full Length (WT) and Tmem178 mutant (L212W/M216W) constructs in pMX-blasticidin retroviral vectors to generate retrovirus used to infect BMDMs. Medium was replaced 24 hours later and the supernatant containing the retrovirus was collected after an additional incubation of 24 hours. BMDMs were infected with 50% supernatant containing indicated retrovirus in α-10 medium containing 10% CMG 14-12 as source of M-CSF and 4ng/ml polybrene for 24 hours. Infected cells were then selected with 1ng/ml blasticidin for 48 hours before being used for indicated experiments.

**IL-1β ELISA.** IL-1β levels were measured using the commercially available ELISA kit (eBioscience, USA) in peritoneal fluids, liver lysates, and/or serum from mice injected with 5 or 10mg/Kg LPS for 6 or 16 hours. For *in vitro* experiments, 0.5 million BMDMs were plated in 6 well plates for 3 days and starved overnight in serum and cytokine free α-MEM media (GIBCO, USA) before stimulation. BMDMs were stimulated with either LPS alone (100ng/ml) or with nigericin (15μM) for indicated time. For NLRP3 inflammasome signaling blockade, BMDMs were treated 1h before adding nigericin with CuET (5μM; Sigma). Medium was collected, centrifuged at 1,500 rpm for 5 min and the supernatant was stored at -80°C until used for ELISA assay according to manufacture procedures.

**Western-blot analysis**. 1 million cells were cultured in 60 $cm^2$ plates for 3 days in α-10 medium with 10% FBS + 10% CMG 14-12 and overnight starved in plain medium before stimulation. BMDMs were stimulated with LPS (100ng/mL) or LPS + nigericin (15μM) for indicated time and then transferred on ice and lysed in RIPA lysis buffer (20 mM Tris-HCl (pH 7.5), 150 mM NaCl,

1mM $Na_2EDTA$, 1mM EGTA, 1% NP-40, 1% sodium deoxycholate, 2.5mM sodium pyrophosphate, 1mM beta-glycerophosphate, 1mM $Na_3VO_4$, 1µg/ml leupeptin) supplemented with protease and phosphatase cocktail inhibitor (GenDEPOT, USA). Protein concentrations were determined by using Bradford assay Reagent (Bio-Rad, USA) and equal amount of protein (50µg) were subjected to SDS-PAGE (10%). The proteins were transferred to PVDF membrane using wet transfer method in transfer buffer (Bio-Rad, USA) supplemented with methanol for 1 hour and 10 min. Blocking was done in 5% BSA for 1 hour before the overnight primary antibody incubation. Primary antibodies included phospho-IκBα (Cell signaling Technology, USA; 2859S; 1:1000) , total IκBα (Cell signaling Technology, USA; 4814S; 1:1000), pro-IL-1β [35 KDa] and cleaved IL-1β [17 KDa] (Abcam, USA; ab234437, 1:1000), NLRP3 (AdipoGen Life Sciences, USA; AG-20B-0014-C100, 1:1000), GSDMD [53 KDa] and cleaved GSDMD [32 KDa] (Abcam, USA; ab209845, 1:1000), pro-caspase-1 (Abcam, USA; ab179515, 1:1000), HA-tag (Cell signaling Technology, USA ; 3724 ; 1:1000), β-actin (Sigma, A5441, 1:5000, MO, USA). The respective HRP- tagged secondary antibodies were incubated for 1 hour before development with luminol based substrate (ThermoFisher Scientific, USA) using ChemiDoc Imaging system (BioRad, USA). The blots were analyzed using ImageJ software.

**Quantitative real-time PCR analysis**. BMDMs (0.5 million) were plated in 6 well plates for 3 days in α-10 medium with 10% FBS + 10% CMG 14-12 and overnight starved in plain medium before stimulation. Stimulation with LPS alone (100ng/ml) or LPS with nigericin (15µM) was done for indicated time. The total RNA was extracted using TRIZOL (Invitrogen, USA) and RNA Mini kit (Qiagen, USA).

Reverse transcription was conducted to generate complementary DNA using high-capacity cDNA Reverse Transcription Kit (Applied Biosystems, CA, USA). Quantitative PCR was performed with

SYBR Green PCR Master Mix (Applied Biosystems) using the 7300 Real Time PCR System (Thermo Fisher Scientific, MA, USA). Relative mRNA expression was calculated by the ΔΔCT method normalizing against cyclophilin B. The sequences of the specific primers were listed:

*Il-1β*, Forward: GCTTCCTTGTGCAAGTGTCTGA

Reverse: TCAAAAGGTGGCATTTCACAGT

*Tmem178*, Forward: ATGACAGGGATATTTTGCACCAT

Reverse: CCGGTTCAAGTCATAGGAGACACT

*Stim1*, Forward: GGCGTGGAAGTCATCAGAAGT

Reverse: TCAGTACAGTCCCTGTCATGG

*Caspase1*: Forward: ACAAGGCACGGGACCTATG

Reverse: TCCCAGTCAGTCCTGGAAATG

*Cyclophilin B*, Forward: ACTGAATGGCTGGATGGCAAG

Reverse: TGCCCGCAAGTCAAAAGAAAT

**Single-cell $Ca^{2+}$ measurements.** WT and *Tmem178*$^{-/-}$ BMDMs were plated in 29mm glass bottom dishes. Adherent cells were incubated with 2 μM Fura-2 diluted in HANKS balanced salt solution (HBSS) plus 2mM $CaCl_2$, 1mM $MgSO_4$ and 10 ng/ml M-CSF for 30 min in dark. Next, cells were washed twice with $Ca^{2+}$ free HBSS containing 1mM MgSO4, and maintained in $Ca^{2+}$ free HBSS buffer plus 10% CMG 14-12. The stained cells were further treated with 15μM nigericin or PBS followed by addition of 2mM $CaCl_2$ during the measurement of calcium fluxes. An Olympus IX-71 inverted microscope with a Lamda-LS illuminator, Fura-2 (340/380) filter set, a 20 × 0.3 N A. objective lens, and a Photometrics Coolsnap HQ2 CCD camera was used to capture images at a frequency of 1 image pair every 2 s. Relative fluorescence ratio at wavelengths of 340nM and 380nM (F340/F380) was utilized for the assessment of basal cytoplasmic calcium level. The calcium fluxes were quantified by the area under the curve, to adjust for differences in basal calcium levels between WT and *Tmem178*$^{-/-}$ BMDMs.

**Mitochondrial functional assays.** Glycolysis and mitochondrial oxidative phosphorylation were estimated by measuring oxygen consumption rate (OCR) using mitochondrial stress test kit (Agilent, USA) and analyzed with Seahorse XFe96 bioanalyzer (Agilent, USA). BMDMs from WT and *Tmem178*$^{-/-}$ mice were seeded at 8x10$^{4}$ cells per well in the Seahorse analysis plates (Agilent, USA) and stimulated 36-48 hours later with PBS (control, n=8/group) or LPS (100ng/mL for 4 hours; n=8/group). Cells were washed and resuspended in xeno-free (XF) media (containing 10mM glucose, 2mM glutamine, and 1mM pyruvate). OCR was measured in response to 1.5μM oligomycin, 2μM fluoro-carbonyl cyanide phenylhydrazone (FCCP), and 1μM rotenone and antimycin A (Agilent, USA). Measurements were taken every 5 minutes. Data generated (Mean and error ± SD) was extrapolated in GraphPad Prism 9.

**Analysis of mitochondrial damage and mitochondrial ROS.** BMDMs (0.5 million) from WT and *Tmem178*$^{-/-}$ mice were plated in 60mm petri dishes for 24 hours in α-10 medium with 10% FBS + 10% CMG 14-12. The cells were replenished with fresh media before stimulating with LPS alone (100ng/ml) for 4 hours or followed by nigericin (15μM) for 45 min. Cells were trypsinized for 5min at 37°C and resuspended in incomplete α-10 medium with 50nM MitoTracker Green (Invitrogen, M7514) or 50nM MitoTracker Deep Red (Invitrogen, M22426) for 15 min at 37°C to detect mitochondrial mass and membrane potential, respectively. Cells were washed with incomplete medium α-10 medium and resuspended in FACS buffer. For mitochondrial ROS (mtROS), cells treated with LPS and/or nigericin were washed with HBBS/Ca/Mg media (GIBCO) and then loaded with 1μM MitoSOX (Invitrogen, M36008) in HBBS/Ca/Mg media for 10 min at

37°C. Cells were washed thrice with HBBS/Ca/Mg and resuspended in FACS buffer. Analysis was performed using the BD-X20 flow cytometer. 10,000 MitoTracker Green and MitoTracker Deep Red positive cells were acquired in the FL1 and FL4 channel, respectively, and MitoSOX positive cells in the FL2 channel. Data analysis was performed using the FlowJo software to calculate the frequency of FL1 positive and FL4-low cells for analysis of damaged mitochondria or FL2 positive cells for mitochondrial ROS (mtROS). In some experiments, cells were treated with 25 or 50µM 2-APB (Sigma) to block SOCE for 30 min before nigericin stimulation and subjected to MitoSOX measurement as described above.

**CuET drug administration.** To block NLRP3 inflammasome, LCMV-infected mice received CuET (1mg/Kg from a 4mg/ml stock dissolved in DMSO; Sigma) intraperitoneally starting two days prior to LCMV infection and continuing every other day for a total of 5 doses. The control mice received vehicle control prepared in variable amount of DMSO diluted in PBS according to the weight of the mice.

**IL-1β neutralizing antibody administration.** To neutralize IL-1β, LCMV-infected mice were given i.p injections of the neutralizing IL-1β monoclonal antibody (10µg; Invivogen) dissolved in sterile water, starting 4 hours after LCMV infection and continuing every other day for a total of 30 days.

**Human sJIA Plasma for macrophage stimulation.** Plasma was obtained from aged matched healthy donors or sJIA patients in the Pediatric Rheumatology Clinic at Lucile Packard Children's hospital and at UCSF enrolled after consent. sJIA patients include both males and females, age 7-18 years, of White, Hispanic/White or Asian heritage, with joint damage and under treatment with *anti*-TNF (Etanercept) alone, or in combination with NSAID, MTX, and prednisone. Healthy donors had similar demographics as the sJIA patients, as described in (16, 18). BMDMs (0.5 million) from WT or *Tmem178*$^{-/-}$ mice were plated in 6 well plates for 3 days in α-10 medium with 10% FBS + 10% CMG 14-12 and overnight starved in plain medium before stimulation. Cells were exposed to 10% plasma from sJIA or healthy patients for 16 hours and further stimulated with LPS (100ng/mL; 4 hours) along with nigericin (7.5μM; 45 minutes). The culture supernatants were collected for analysis of IL-1β levels by ELISA and total RNA was extracted using TRIZOL (Invitrogen, USA) and RNA Mini kit (Qiagen, USA). THP-1 cells (ATCC®TIB-202TM) were obtained from ATCC, USA and maintained in RPMI-1640 medium supplemented with 10% FBS. 0.5 million THP-1 cells in 6 well plates were stimulated with 100nM PMA for 48 hours to allow macrophage differentiation. Cells were then exposed to 10% plasma from sJIA or healthy patients for 12 hours followed by stimulation with LPS (100ng/mL; 4 hours) and nigericin (7.5μM; 45 minutes). Total RNA was extracted using TRIZOL (Invitrogen, USA) and RNA Mini kit (Qiagen, USA). The samples were stored in -80$^{0}$C until processing.

**Principal component, differential expression analyses, and heatmaps.** Data from 160759 genes (GSE147608) obtained from monocytes in active sJIA patients or healthy controls were compared to each other to derive the differentially expressed genes ($p<0.05$). Genes with zero counts across all the samples were excluded for further analysis. Next, the DESeq2 R package was used for differential gene expression analysis. DESeq method does internal normalization by dividing counts for each gene by the geometric mean (calculated for each gene across samples). Volcano plots showing the relationship between log2 fold change and significance (-log 10 * p-value) were constructed using the ggplot2 R package. The volcano plot includes differentially expressed genes with the threshold of an unadjusted ($\text{p-value}_{\text{unadjusted}} < 0.05$) and adjusted (19) $\text{p-value}_{\text{adjusted(BH)}} <$

0.05). Number of differentially expressed genes detected using the DESeq2 method are shown below.

| **Unadjusted p-value** | | **Adjusted p-value (BH) Benjamini & Hochberg (1995)** | |
|---|---|---|---|
| UP-regulated | Down-regulated | UP-regulated | Down-regulated |
| 11097 | 18925 | 8295 | 14534 |

### Supplementary Figures and Legends

### Supplementary Fig 1

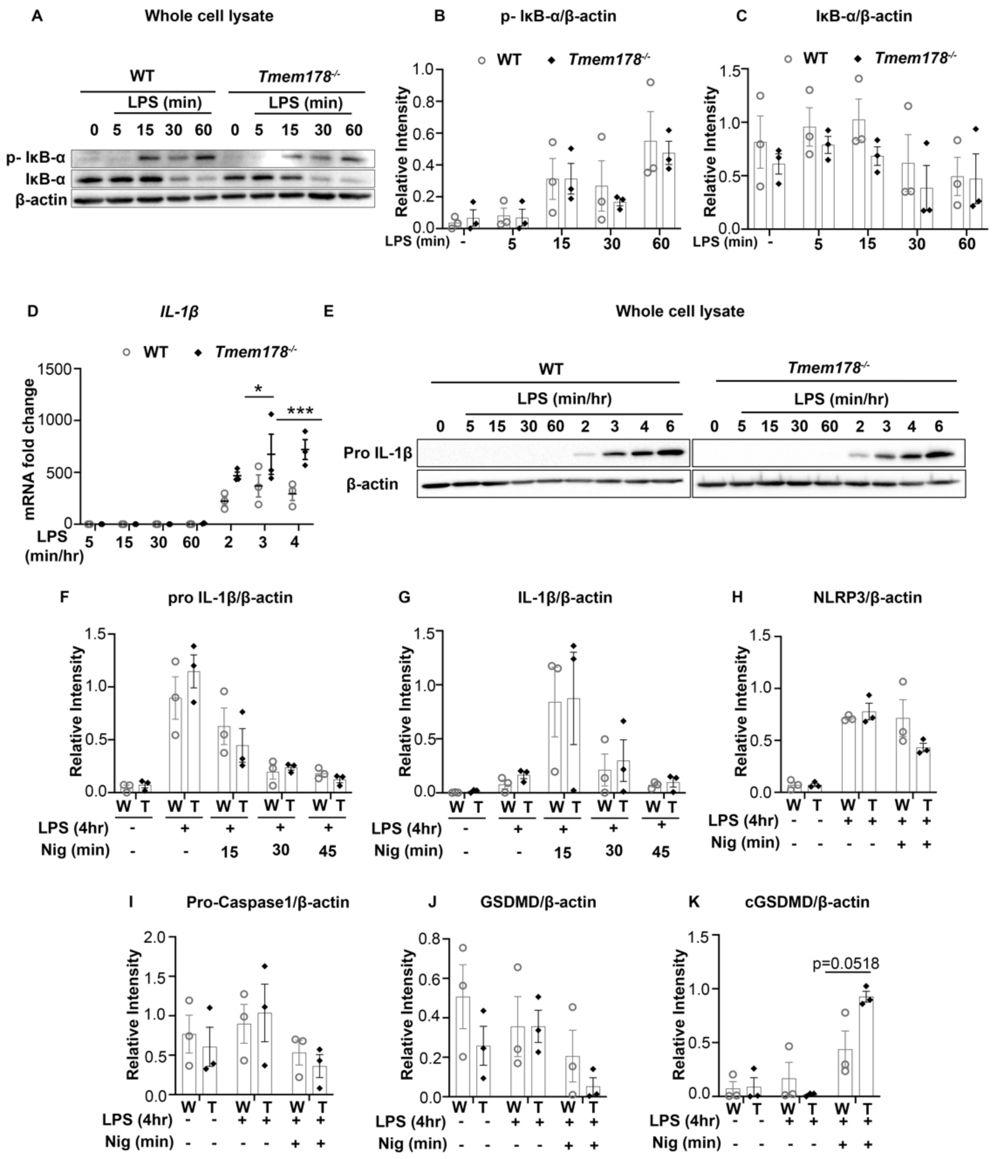


**Supplementary Fig 1.** Role of Tmem178 in NLRP3 inflammasome priming. (A-C) BMDMs from WT and *Tmem178*$^{-/-}$ mice were primed with 100ng/ml LPS for indicated time to detect abundance of phosphorylated and total IκB-α proteins (A). Densitometric analysis of phosphorylated IκB-α (B) and IκB-α (C) was performed on three different sets of experiments. (D-E) Analysis of *IL-1β* transcripts and pro-IL-1β protein levels in WT and *Tmem178*$^{-/-}$ stimulated with 100ng/ml LPS for indicated time. (F-K) Densitometric analysis of BMDMs primed with 100ng/ml LPS for 4 hr followed by nigericin (15μM) for 45 min for (F) pro-IL-1β, (G) mature IL-1β, (H) NLRP3, (I) pro-caspase-1, (J) GSDMD, (K) cleaved GSDMD proteins in WT (W) and *Tmem178*$^{-/-}$ (T) lysates. Data are expressed as mean ± SEM (n = 3/group). Statistical significance is determined by two-way ANOVA for multiple comparisons (D) or two-tailed t-test (K). $*P < 0.05$ and $***P < 0.001$.

**Supplementary Fig 2**

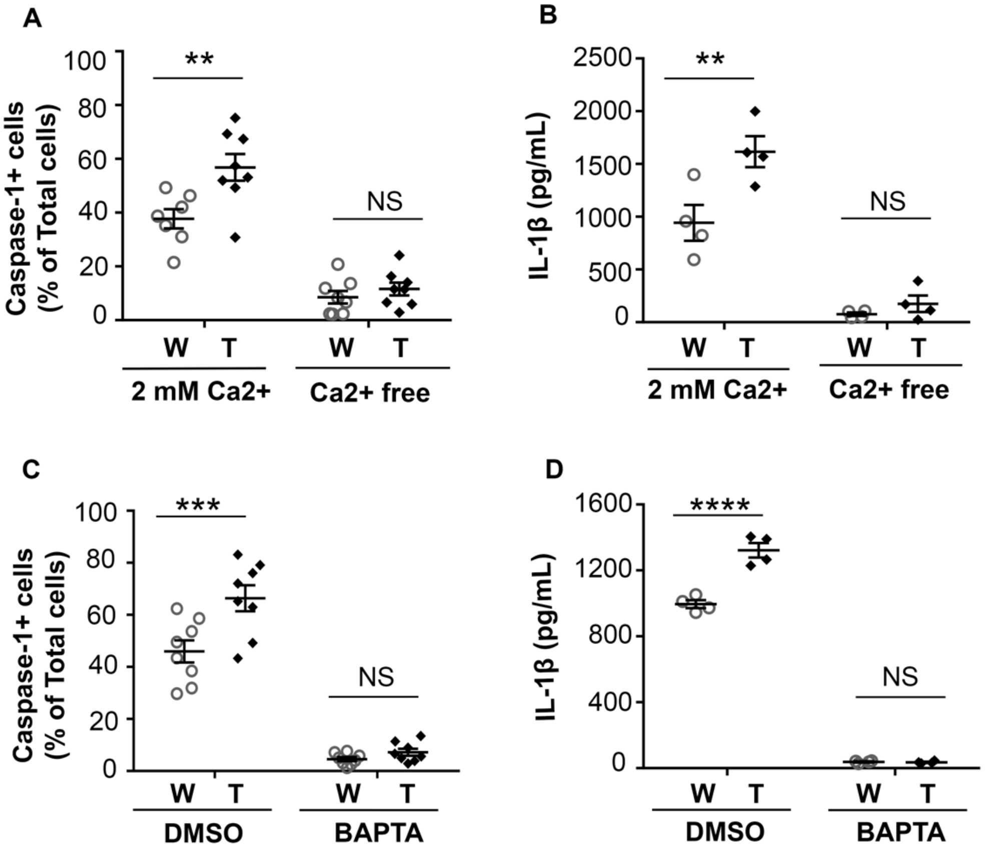


**Supplementary Fig 2.** Role of cytosolic $Ca^{2+}$ in NLRP3 inflammasome activation. (A-B) BMDMs primed with LPS (100ng/ml, 4hr) in complete media were switched to $Ca^{2+}$ free-HBSS or 2mM $Ca^{2+}$-HBSS followed by nigericin stimulation (15µM, 45min). (A) Quantification of cells with active caspase-1 (n= 7/8). (B) Mature IL-1β levels in culture supernatants (n=4). (C-D) BMDMs primed with LPS (100ng/ml, 4 hr) and treated with DMSO or 10µM BAPTA-AM for 30 min prior to nigericin (15µM, 45min) stimulation. (C) Percentage of cells with active caspase-1 (n = 8) and (D) IL-1β levels in culture supernatants. Data are presented as mean ± SEM. Statistical significance is determined by two-way ANOVA for multiple comparisons. $^{**}P < 0.01$ , $^{***}P < 0.001$ and $^{****}P < 0.0001$.

**Supplementary Fig 3**

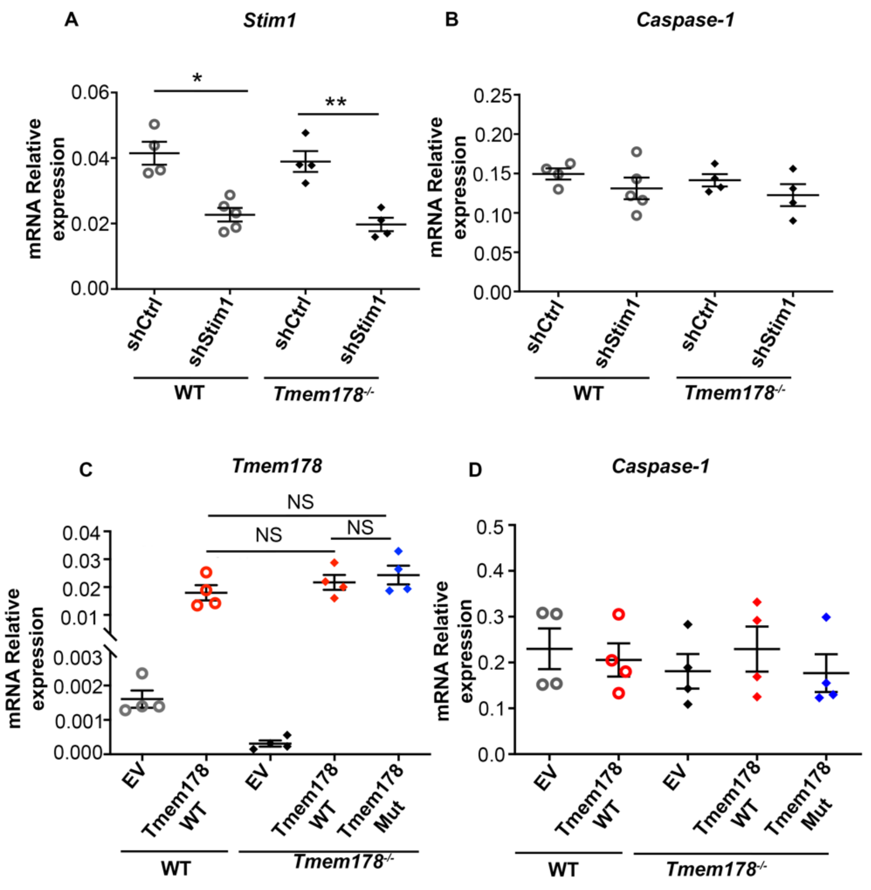


**Supplementary Fig 3.** mRNA levels of *Stim1*, *caspase-1* and *Tmem178*. (A-B) WT and *Tmem178*$^{-/-}$ BMDMs expressing shRNA control or shRNA Stim1 were subjected to RT-PCR analysis to measure *Stim1* (A) and *caspase-1* (B) transcripts. (C-D) WT or *Tmem178*$^{-/-}$ BMDMs expressing empty vector (EV), HA-tagged full length Tmem178 (Tmem178-WT) or the mutant lacking Stim1 binding site (Tmem178-Mut) were subjected to RT-PCR analysis to measure (C) *Tmem178* and (D) *caspase-1* transcripts (n=4/group). Data are presented as mean ± SEM. Statistical significance was determined by two-way ANOVA for multiple comparisons. *$P < 0.05$, **$P < 0.01$.

**Supplementary Fig 4**

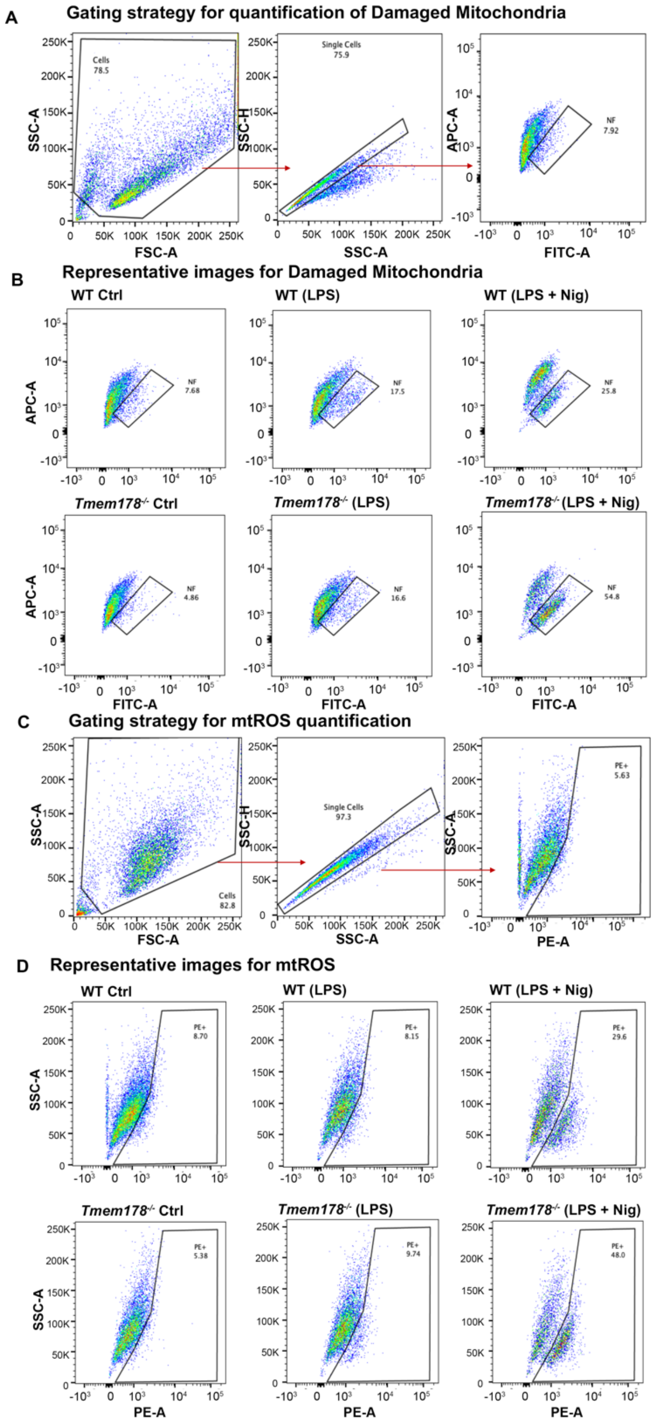


**Supplementary Fig 4.** Loss of Tmem178 induces mitochondria dysfunction. Gating strategy (A and C) and representative images of (B) damaged mitochondria (MitoTracker Green$^{high}$ and

MitoTracker Red$^{low}$) and (D) mtROS (MitoSOX$^{high}$) from WT and *Tmem178*$^{-/-}$ BMDMs primed with 100ng/ml LPS for 4hr followed by nigericin treatment (15μM, 45 min).

**Supplementary Fig 5**

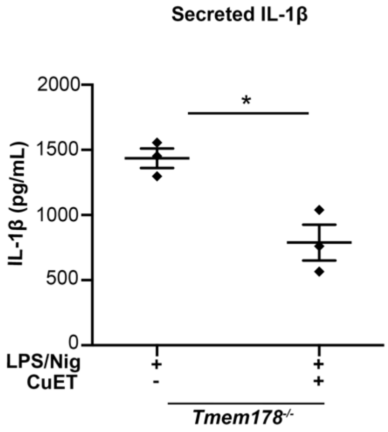


**Supplementary Fig 5.** Reduced IL-1β release by inflammasome inhibitor. IL-1β levels in culture supernatants measured by ELISA in *Tmem178*$^{-/-}$ BMDM primed with LPS (100ng/ml, 4hr) followed by CuET treatment (5µM) or vector control for 1hr and stimulation with nigericin (15µM, 45 min) (n=3/group). Data are presented as mean ± SEM. Statistical significance is determined by two-tailed *t*-test. *$P < 0.05$.
